# Analysis of the molecular determinants for furin cleavage of the spike protein S1/S2 site in defined strains of the prototype coronavirus murine hepatitis virus (MHV)

**DOI:** 10.1101/2023.01.11.523687

**Authors:** Annette Choi, Ekaterina D. Kots, Deanndria T. Singleton, Harel A. Weinstein, Gary R. Whittaker

**Affiliations:** Departments of Microbiology & Immunology, Cornell University, Ithaca, NY, USA; Department of Physiology & Biophysics, Weill Cornell Medicine, New York, NY, USA; Institute for Computational Biomedicine, Weill Cornell Medicine, New York, NY, USA; Public & Ecosystem Health, Cornell University, Ithaca, NY, USA

## Abstract

We have analyzed the spike protein S1/S2 cleavage site of selected strains of MHV by the cellular protease furin, in order to understand the structural requirements underlying the sequence selectivity of the scissile segment. The probability of cleavage of the various MHV strains was first evaluated from furin cleavage scores predicted by the ProP computer software, and then cleavage was measured experimentally with a fluorogenic peptide cleavage assay consisting of S1/S2 peptide mimics and purified furin. We found that *in vitro* cleavability varied across MHV strains in line with predicted results—but with the notable exception of MHV-A59, which was not cleaved despite a high score predicted for its sequence. Using the known X-Ray structure of furin in complex with a substrate-like inhibitor as an initial structural reference, we carried out molecular dynamics (MD) simulations to learn the modes of binding of the peptides in the furin active site, and the suitability of the complex for initiation of the enzymatic cleavage. We thus identified the 3D structural requirements of the furin active site configuration that enable bound peptides to undergo cleavage, and the way in which the various strains tested experimentally are fulfilling these requirements. We find that despite some flexibility in the organization of the peptide bound to the active site of the enzyme, the presence of a histidine at P2 of MHV-A59 fails to properly orient the sidechain of His194 of the furin catalytic triad and therefore produces a distortion that renders the peptide/complex structural configuration in the active site incompatible with requirements for cleavage initiation. The Ser/Thr in P1 of MHV-2 and MHV-S has a similar effect of distorting the conformation of the furin active site residues produced by the elimination of the canonical salt-bridge formed by arginine in P1 position. This work informs a study of coronavirus infection and pathogenesis with respect to the function of the viral spike protein, and suggests an important process of viral adaptation and evolution within the spike S1/S2 structural loop.

## Introduction

The *Coronaviridae* family of viruses includes highly diverse members that emerge from a wide range of animal reservoirs, from bats and birds to mammals and rodents (1, 2). A primary determinant of tropism and transmission is the viral spike glycoprotein (S) that is composed of two domains - S1 (responsible for receptor binding) and S2 (mediating membrane fusion) (3). Many members of the *Coronaviridae* contain a distinct protease cleavage-activation site at the S1/S2 interface of S, which meets the consensus motif for cleavage by the ubiquitous protease furin composed of arginine (R) residues required at the core P4 and P1 cleavage residues (R-x-x-R) and a preference for a basic residue at P2 (*i.e*., R-x-K/R-R) (4). In many coronaviruses, such furin cleavage sites are thought to regulate S protein function, including “priming” the process of fusion-activation. Such sites can be found in human, avian, and rodent coronaviruses, including across the betacoronavirus genus, but are not typically present in SARS-like sarbecoviruses (including in bat reservoirs). A notable exception is SARS-CoV-2 which contains a furin cleavage site that is atypical in terms of the positioning of the basic (R) residues (R-R-A-R), and which has evolved over the course of the COVID-19 pandemic to include additional basic residues and/or histidine residues (5, 6).

Murine hepatitis virus (MHV) is a betacoronavirus (sub-genus embecovirus) often used as a model to study pathogenic mechanisms of coronavirus biology. MHV exists as distinct strains that differ considerably in their virulence and carry different organotropisms (7). For example, MHV-1 and MHV-S are less virulent compared to strains MHV-2, MHV-3, and MHV-A59; while MHV-JHM is neurotropic. These strains represent distinct laboratory-isolated variants that are distinct pathogenically, but highly homologous in their overall S sequences; see Figure 1 which depicts phylogenetic relationships across selected coronaviruses.

**Figure 1.**
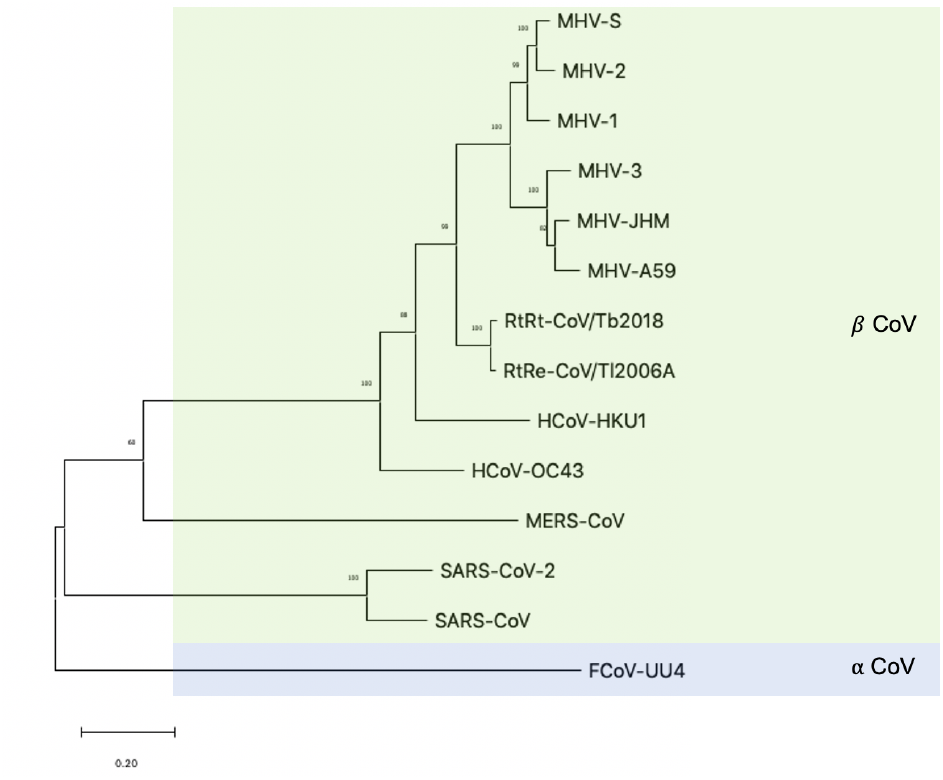
Phylogenetic tree of selected coronaviruses. Phylogenetic tree of spike protein sequences. The tree was constructed using MegaX, 100 bootstrap replicates from a spike alignment. See Methods for accession numbers used.

As is common in the embecoviruses, and similar to SARS-CoV-2, the S protein of many MHV strains contains an RXXR sequence within the S protein, suggesting that they contain a functional furin cleavage site and are cleaved by furin. Here, we examined cleavage by furin of six representative strains of MHV that have a range of pathogenic outcomes following initial respiratory infection, which includes pneumonitis, fulminant hepatitis, and encephalitis. The tested strains differ in the positions at which His or Ser/Thr residues appear in key protease cleavage positions.

## Results

### In vitro studies of the modes of interaction of the MHV strain peptides with the furin active site

To predict furin cleavage probabilities, we first utilized the ProP computer software (https://services.healthtech.dtu.dk/service.php?ProP-1.0); see ref (8), which predicts cleavage by proprotein convertases (including furin) using artificial neural networks. A score of > 0.5 is considered the default threshold for cleavage (Dukert et al. (2004) (Table 1). The ProP scores predicted that furin cleavage should occur for strains MHV-1, MHV-JHM/MHV-3, and MHV-A59, with furin cleavage minimal or absent for MHV-2 and MHV-S… Figure 1 shows the predicted ProP scores for MHV-1 (0.84), MHV-2, (0.21), MHV-JHM/3 (0.87), MHV-S (0.10), and MHV-A59 (0.77), suggesting that only MHV-2 and MHV-S would fail to be cleaved by furin.

**Table 1.**
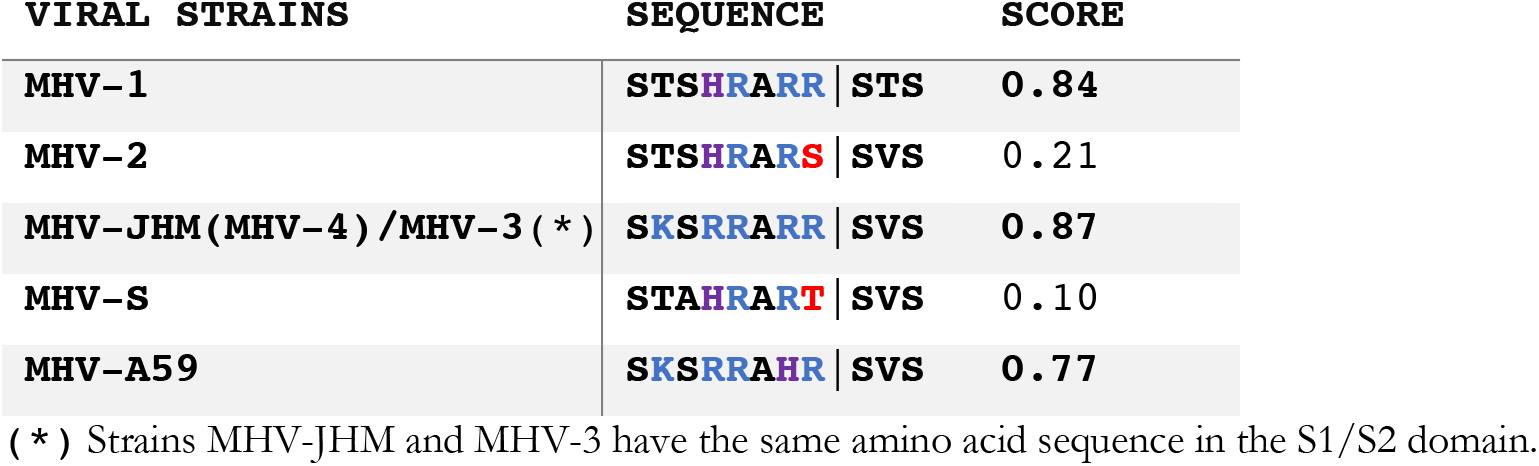
Sequences of MHV strains tested and ProP scores for furin cleavage probability.

These predictions were then compared to the results from determinations of cleavability in experiments using a fluorogenic peptide cleavage assay shown in Figure 2. Notably, although MHV-A59 was predicted by the ProP scores to be cleaved by furin, it is found to be only minimally processed in our biochemical peptide cleavage assay.

**Figure 2.**
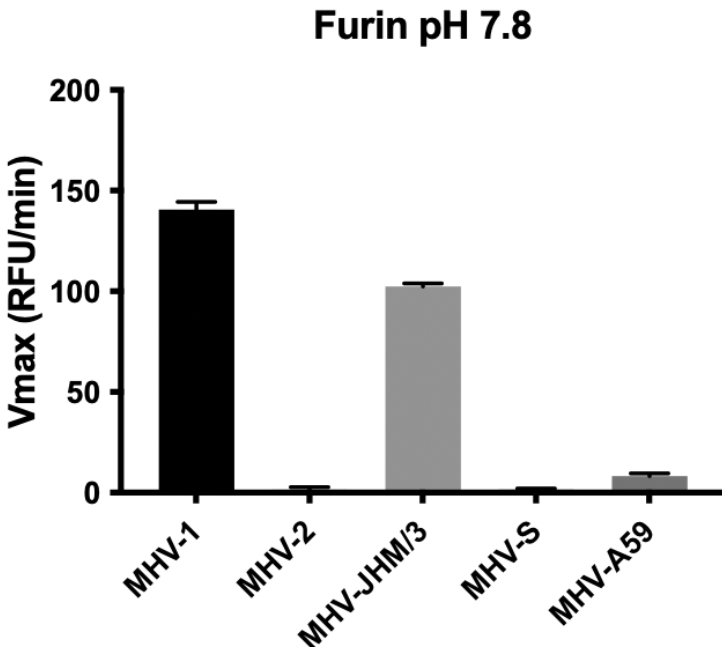
Processivity of Furin Cleavage at pH 7.8 on MHV S1/S2 Peptide Mimics. The figure shows the Vmax of cleavage by furin at pH 7.8 when mixed in buffer with select MHV S1/S2 peptide mimics. *Strains MHV-JHM and MHV-3 have the same amino acid sequence in the S1/S2 domain

As previous studies show that the presence of defined histidine residues within the glycoproteins of viruses can affect the conformational changes the glycoprotein undergoes during viral entry in a pH-dependent manner (Kalani et al., 2013), we further assessed furin cleavage at different pH values to investigate whether the pH would affect the processivity of furin cleavage based to the presence of the P2 histidine. Thus, considering that our selected MHV strains contain furin cleavage sites with a neighboring upstream histidine residue, this led us to test furin cleavage at pH values. Fig. 3 shows the Vmax for furin cleavage of the MHV strains at different pH values, specifically, pH 6.8. pH 6.3, and pH 5.5.

**Figure 3.**
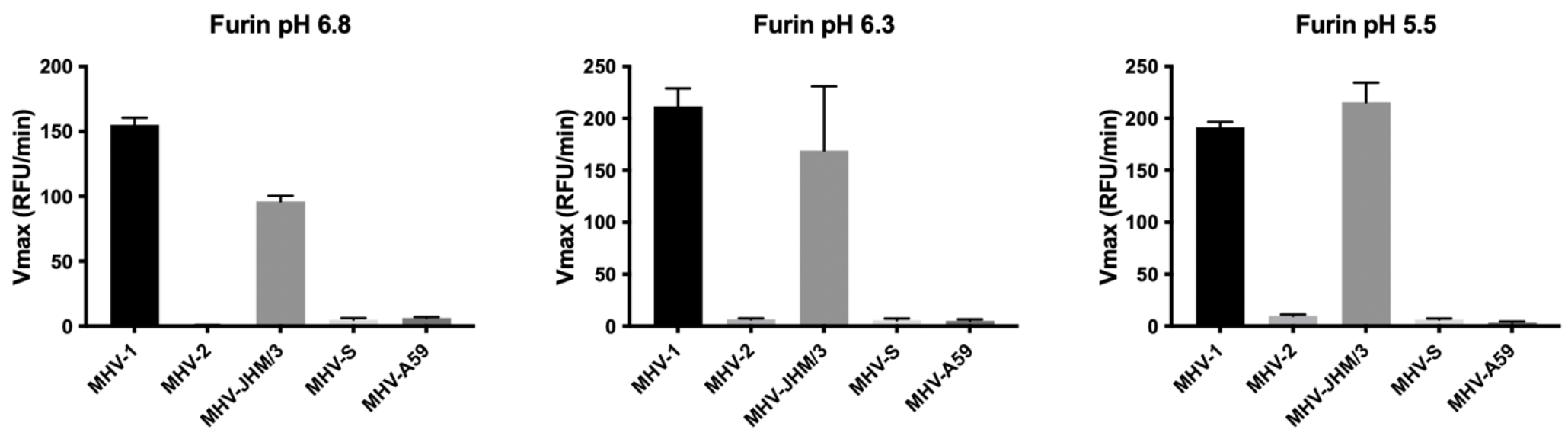
Processivity of Furin Cleavage at pH 6.8,6.3 and 5.5 on CoV Peptide Mimics. The table above depicts the velocity of furin protease at pH 6.8,6.3, and 5.5 when mixed in buffer with select CoV linear peptide mimics. *Strains MHV-JHM and MHV-3 have the same amino acid sequence in the S1/S2 domain

As outlined in Jaimes et al. (2019), the set-up of the fluorogenic peptide cleavage assay, precludes a comparison of cleavability of the protease across strain and pH values, but we can compare cleavability between strains within the same pH value. The trend observable for specific MHV strains across pH values is that, as predicted with ProP, the MHV-1 and MHV-JHM/3 strains showed consistent cleavage by furin irrespective of pH, while MHV-2 and MHV-S showed minimal cleavage. These results were expected, especially because MHV-1 and MHV-JHM/3 contain the specific series of amino acid residues recognized as the FCS. However, the cleavage of MHV-A59 was minimal at all pH values tested despite the predicted score of 0.77 in the ProP analysis.

### Computational studies of the modes of interaction of the MHV strain peptides with the furin active site

#### Structural requirements for the enzymatic cleavage reaction in the furin active site

As a member of serine protease protein family, furin mediates peptide bond cleavage using a catalytic triad composed of residues Asp153, His194, and Ser368 (Figure 4) (4, 9). The nucleophilic attack on the carbonyl carbon of the substrate residue in the P1 position is carried out in the first step of the hydrolysis mechanism by the Ser368 oxygen. The Nε-atom of His194 acts as a proton switch, first by receiving a proton from the Ser368 hydroxyl group and transferring it to the N-terminus group of P1’ peptide segment which is released when the P1-P1’ peptide bond is broken. Ser368 forms a covalent complex with the P1 segment (Ser368-O-C-P1-). During the second step, when the activated water molecule performs a nucleophilic attack on the carbonyl atom of the covalent complex, the covalent bond between hydroxyl atom of Ser368 and P1 carbonyl carbon is broken and P1 is released, and the His194 again acts as a proton switch and delivers it to the -OH group of Ser368. The steric requirements that this mechanism imposes on the spatial organization of the active site residues includes the following three conditions: 1) The distance between Nε-atom of His194 and hydroxyl oxygen of Ser368 should be less than ~3.3 Å to allow for the proton transfer; 2) The distance between carbonyl atom of the P1 peptide residue and the hydroxyl oxygen of Ser368 should not exceed 3.5 Å to enable the nucleophilic attack of Ser368 on the peptide bond; and lastly, 3) The region around the carbonyl atom of P1 should be accessible to water to enable the cleavage of the covalent complex formed between Ser368 and P1 segment of the peptide during the first step of the cleavage. Therefore, in order to evaluate the feasibility of the enzymatic reaction of each of the investigated peptides representing putative substrates, we monitored these metrics throughout the sampling of the dynamic behavior of the furin-peptide complexes in the MD simulation trajectory results.

**Figure 4.**
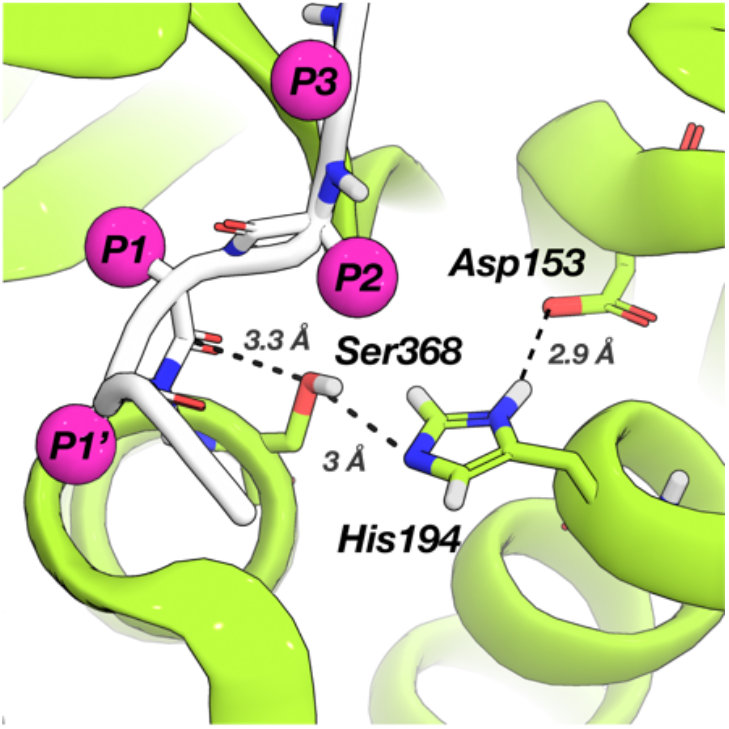
Spatial organization of the furin catalytic triad composed of Asp153/His194/Ser368 in the results of MD simulation of the furin-MHV-3 complex. The orientations of the sidechains of P1-P3 and P1’ residues of the peptide are indicated by the magenta-colored balls.

Specifically, we evaluated for each complex the probability (calculated as the ratio of microstates in which a conformation prevailed) for the orientations of His194, Ser368 and the backbone of the P1 residue of a particular peptide bound in the active site to satisfy the requirements for cleavage (Table 2).

**Table 2.**
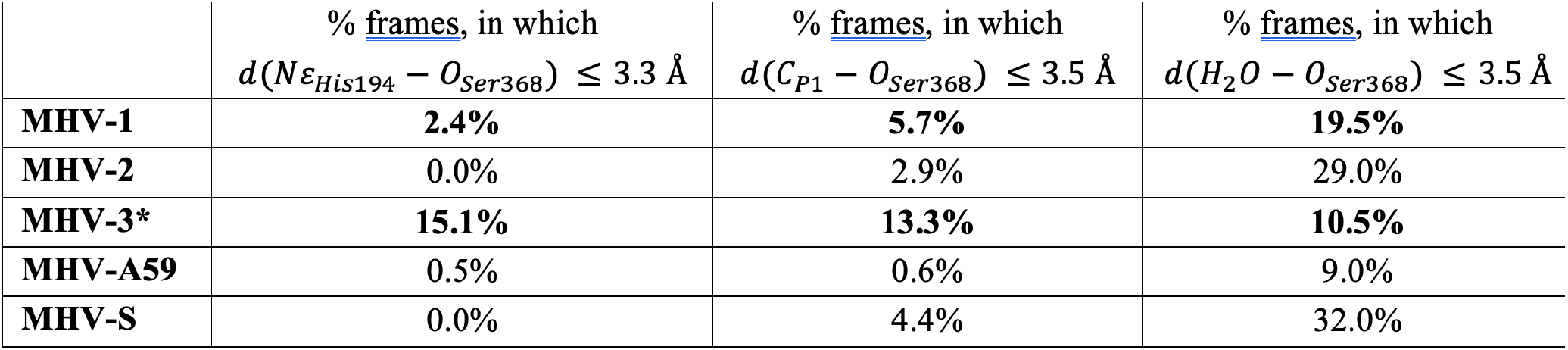
The probability that the furin complex with each of the five MHV strains can initiate the cleavage process satisfies, expressed as the percentage of MD simulation frames in which the criteria imposed on the geometry of the catalytic site residues are satisfied.

The substrates with different sequences are expected to adopt different conformations and interact with different residues in the environment of the active site, therefore they should affect differently the conformation of the active site pocket and produce different geometries of the catalytic site in the resulting complexes. Consequently, the metrics of enzyme residue positions with respect to peptide sites will differ for each peptide. This is clearly seen in Table 2. The quantified occurrences of satisfied criteria show that the strictest geometrical requirement is the formation of the hydrogen bond between sidechains of His194 and Ser368, as in this set of peptides only the complexes with the MHV-1 and MHV-3 strains exhibit occupancies higher than 1%. For the complexes of these two strains, the occurrence of the Ser368 and carbonyl carbon of P1 residue being positioned at a distance suitable for nucleophilic attack is also the highest among the MHV strains tested. This result validates both the criteria and the observations from the MD simulations since the sequences within the S1/S2 domain region of both of these strains are -R-A-R-R- (Figure 2), which is a canonical -P1-to-P4- sequence in furin substrates.

#### MD simulations of the modes of binding of the MHV peptides that determine the cleavage probability

As shown in Table 2, the probabilities of favorable orientations of the P1 backbone towards the furin Ser368, His194 and the complex with MHV-1 are significantly lower than in the MHV-3 complex. From the MD simulations we learned how the differences in the occurrence of catalytic site conformation that are suitable for cleavage are attributable to the difference in the sequences of the two strains. Specifically, the MHV-1 and MHV-3 sequences differ in two positions: P5 (His vs Arg, respectively) and P2’ (Thr vs Val). In the MD trajectories of the furin/MHV-3 complex, the P5 Arg forms a stable salt-bridge with either Glu230 or Glu257. In contrast, the P5 His of MHV-1 forms a hydrogen bond only occasionally, with the backbone of Val231 (Figure S1C-D). The stable sidechain interactions of the MHV-3 P5 residue restricts the flexibility of the peptide in the groove (Figure S1A). The decreased mobility of the backbone of the peptide in the groove (Figure S1B) will increase the probability of catalytic site conformations favorable for the cleavage. Notably, however, these MD simulations reflect only the initiation state of the catalytic reaction and thus do not reflect all the factors affecting a measured Vmax. Indeed, cleavage assays at neutral pH measure only a slight difference in the Vmax of MHV-1 and MHV-3 cleavage by furin. Therefore, we can assume that initiation of the cleavage reaction is possible if MD ensembles of furin/MHV strain complexes exhibit significant occurrences of the active site conformation required to initiate the cleavage reaction. Here, we find that this requirement is met only for the MHV-1 and MHV-3 strains. In the simulations of complexes of furin with substrates presenting a P1 arginine (as in the canonical P1-to-P4 sequence R-A-R-R) the P1 residue is positioned in close vicinity to the calcium binding site and forms salt-bridges with Asp258 and Asp306 (4, 10). Indeed, we observe this positioning of the P1 arginine in MD simulation of the furin complexes MHV-1 and MHV-3 strains (Figure 5a, d). These interactions act as anchors and hold P1 residue in place, at the same time positioning its backbone in the vicinity of Ser368, which sets up the nucleophilic attack. For the MHV-2 and MHV-S complexes, however, we find that His194 fails to form a hydrogen bond with Ser368 (Table 2), which precludes the first stage of cleavage process for these MHV strains. In both these strains the P1 substrate position contains a polar amino acid (serine and threonine in MHV-2 and MHV-S, respectively) with neutral charge and shorter sidechain than that of the canonical arginine. Consequently, the salt-bridge interactions at ~3.5 Å with Asp258 and Asp306, are absent in the complexes of furin with MHV-2 and MHV-S (Figure 45b, e). The corresponding minimal sidechain distances are therefore above 6 Å (Figure 5f, left), so that the displacement of the backbone moves the P1 residue away from the hydroxyl oxygen of Ser368 (Table 2, Figure 5b, e) and distorts the overall geometry of the catalytic site making cleavage impossible.

**Figure 5.**
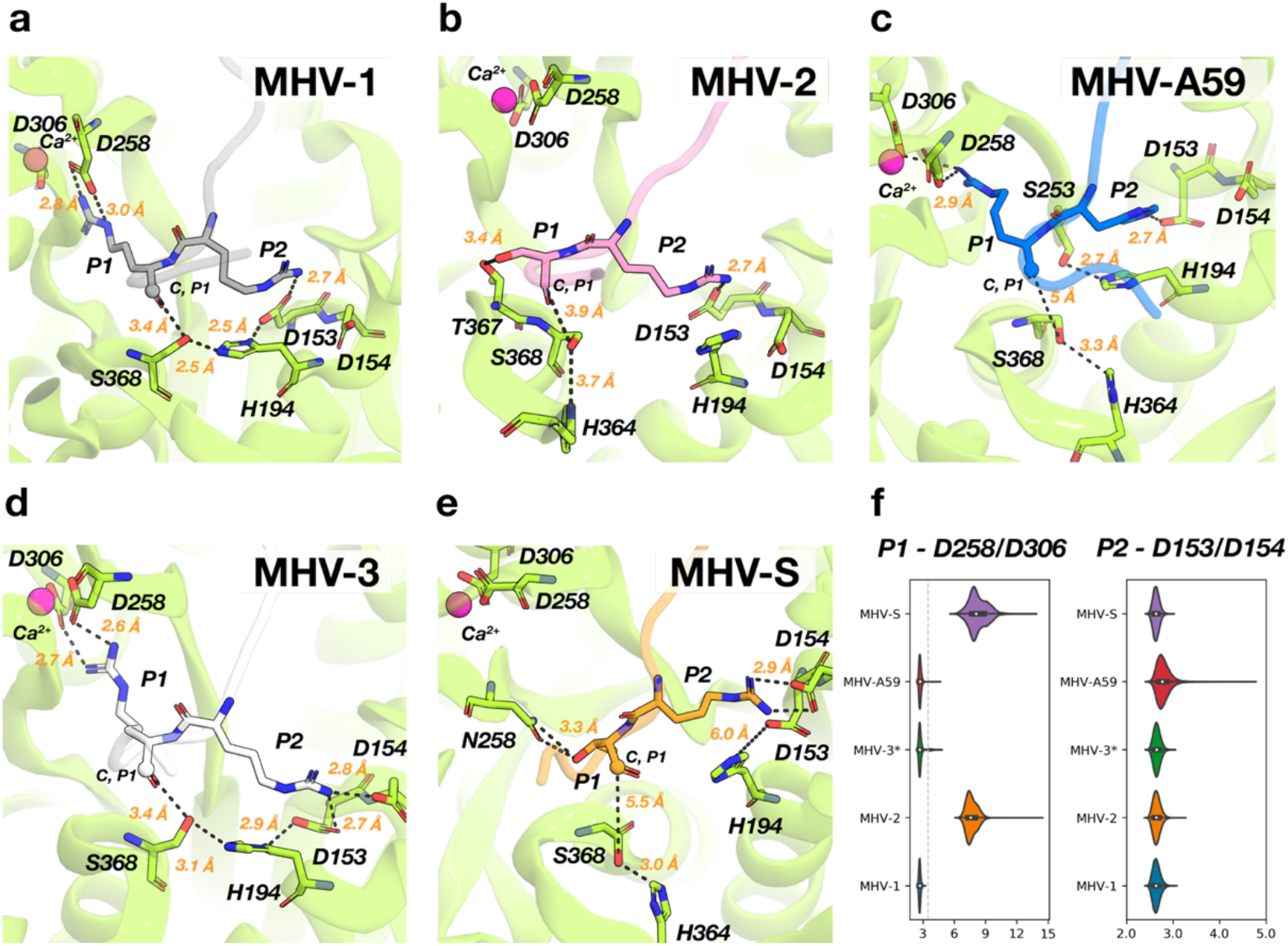
A-E) Representative conformations of active site region of furin in complex with MHV-1, -2, -A59, -3 and -S strains from the MD simulations. F) Distribution of the minimum distances between sidechain atoms of P1 residue and Asp258/As306 (left) and P2 residue and Asp153/Asp154 (right) computed from the MD simulation of furin complexes with the five MHV strains.

A similar distortion of the active site configuration needed for cleavage occurs when the substrate P2 position is not arginine. Thus, the canonical P2 arginine in all MHV strains tested – except MHV-A59 – is engaged in salt-bridges with Asp153 or/and Asp154 of the furin catalytic site (Figure 5a-d). When the P2 position is occupied by arginine, the observed minimum sidechain distance measured between the nitrogen atoms of the P2 residue and oxygens of Asp153 and Asp154 are <3 Å (Figure 5f, right). These salt-bridges assist in keeping the backbone of P1 and P2 residues in place and help to orient His194, so that it can act as a proton switch during as described (Figure 4, 5a, d).

In MHV-A59, where the P2 position is occupied by histidine (Figure 2) which has a shorter sidechain and doesn’t a carry positive charge at neutral pH, we observe the formation of the hydrogen bond between P2 histidine and Asp153/Asp154 sidechains, (Figure 5f, right), but unlike arginine, histidine in P2 position fails to orient His194 appropriately (Figure 5c). This disrupts the spatial organization of the catalytic triad and prevents the proton transfer from Ser368 to His194 that is required to initiate the cleavage. Moreover, the way in which the bound MHV-A59 peptide is positioned in the active site of furin is not compatible with the steric requirements for the cleavage reaction. Thus, the distance for nucle ophilic attack interaction between the carbonyl carbon of the MHV-A59 P1 residue and the hydroxyl oxygen of Ser368 is too long (Figure 5c) and almost never falls below 3.5 Å (Table 2). Therefore, conformations that would favor the initiation of cleavage were observed in less than 1% of the MD frames.

## Discussion

Our work advances the understanding of furin-mediated S protein cleavage for the model coronavirus MHV, with the finding that MHV-A59 is not cleaved by furin despite its high predictive score by ProP and the proposal of a structure-based mechanism for this observation. MHV-A59 is a “prototype” MHV/embecovirus commonly used for laboratory studies; however, several MHV-A59 sequences are present in databases, and all MHV stains are likely laboratory adaptations. As such, it is hard to determine a “wildtype” sequence for this family of viruses. Such a sequence would need to be derived from samples taken directly from a rodent. Several candidates for wildtype embecoviruses do exist, including RtRt-CoV/Tb2018 (GenBank: QIM73854.1) (11), with an S1/S2 sequence of **SKARRARR**|**SVS** and a ProP score reported as 0.87 and discussed in ref (2). With only limited sequences available, we consider that MHV-3 may comprise the most appropriate “wildtype” laboratory MHV/embecovirus strain, with a cleaved spike protein to match that expected in wild mice; see ref (12).

We studied six commonly used MHV strains, including MHV-A59, but it is important to note that several studies involving long term passage of MHV-A59 have selected viruses with modified S1/S2 cleavage site that preserve the P2 His residue but introduce residues that would classically be expected to be refractory to furin cleavage (substitutions shown in red in the text below). For instance MHV-ExoN(-) was selected after 250 passages of WT (A59)-MHV in DBT cells and was reported to have reduced syncytium formation (13) and contains a spike S1/S2 sequence of **SKSRGAHR**|**SVS** (GenBank: ATN37896.1). Another example is the selection of MHV/BHK (spike S1/S2 sequence of **SKSRRAHR**|**LGS**) from MHV-A59 following 600 passages in cell culture—in this case also selecting heparin binding activity though a second site on spike (14); however, the authors do not directly comment of fusion activity in this study. In addition, selection of MHV-A59 after 16 weeks of culture in glial cell culture selected a spike S1/S2 sequence of **SKSRRADR**|**SVS** (15), also with minimal levels of cell-cell fusion. Overall, these data suggest that introduction of a His residue into the FCS may represent an early adaption in the process of virus evolution to new hosts, tissue or cells— with additional mutations (i.e., Ser, Thr, Gly, Asp) being introduced later, to more completely inhibit furin-mediated cleavage of the spike protein. It is interesting to note that SARS-CoV-2 has also introduced His residues into its FCS (from PRRAR| S in WA/1 to HRRAR| S in the Alpha and Omicron VOCs, whereas the highly fusogenic and transmissible Delta VOC introduced an Arg residue RRRAR| S) (6, 16). It remains to be determined the effect of introduction of a His residue into the P5 position of the FCS (as seen in MHV-1, MHV-2 and MHV-S) represents in term of adaption in the process of SARS-CoV-2 evolution to the human population. It is notable that no changes have occurred in the second (fusion peptide-proximal) S2’ cleavage site, for either MHV or SARS-CoV-2. Finally, experimental validation of the effect of His substitutions must be carefully evaluated with respect to cell-cell fusion events (syncytial formation) vs. virus-cell fusion events occurring during virus entry and the cell surface or within endosomal compartments.

MHV-2 is the only MHV strain that is widely documented to be “uncleaved’, and not to induce cell-cell fusion; it enters cells via the cathepsin pathway (17). However, cleavability by furin does not seem to be correlated with the relative virulence of the MHV strains. Thus, with respect to the S protein, MHV-S and MHV-A59 are considered more virulent and were cleaved at a sub-threshold by furin compared to MHV-1, which was moderately cleaved by furin and is considered less virulent. These results may suggest that other aspects of the viral genome may be responsible for the increased virulence of MHV-S compared to MHV-A59 which is found to not be cleaved by furin. Within the S protein of MHV-A59, the FCS contains a RAHR| sequence rather than the RARR| sequence seen in MHV-1 and MHV-JHM/3, and the expectation was for. a moderately predicted furin cleavability. Initially, we believed that the histidine was inhibiting furin’s ability to recognize the FCS for MHV-A59 due to the “histidine switch”, based on finding with the conformational changes required for fusion in other virus systems.

One notable finding of our study is the lack of cleavage of the MHV-A59 FCS despite its high ProP score. MHV-A59 has been reported previously to be uncleaved in certain cell lines; i.e., in primary hepatocyte and mixed glial cells culture (as opposed to continuously cultured cells lines) (18). Absent structural information about the cleavage mechanism for this variant, the lack of cleavage could have been attributed to other structural or post-translational modifications elsewhere on the spike – possibly proximal or distal glycosylation (O-linked or N-linked) that is well known to influence cleavage-activation of viral glycoproteins (19, 20). Differences in MHV-A59 spike cleavage might also have been influenced by cell-type specific expression of other (non-furin) members of the proprotein convertase family, which are less stringent with respect the amino acid sequence of the protease substrate (21). Our data clearly show that lack of cleavage of MHV-A59 spike by furin is an inherent property of its **SKSRRAHR**|**SVS** cleavage site, specifically based on the presence of a P2 His residue that is causing a rearrangement of the active site-peptide complex to a non-productive configuration. Thus, the application of the MD simulations in this study has revealed the equilibrium structural configurations of the pre-cleavage active sites of the furin in complex with each of the MHV-strains examined, from which a specific mechanistic hypothesis emerged.

The models obtained from the MD simulations for the complexes of the studied peptides with the furin active site illuminate the structural considerations underlying the probability of cleavage initiation. These considerations are based on the probability of a particular sequence to adopt a conformation in the active site that enables cleavage. To activate the proteolysis mechanism, such a conformation needs to permit the furin catalytic triad residues (Asp153, His194, Ser368) to adopt a proper orientation towards the backbone of the substrate’s P1 residue so as to enable the nucleophilic attack of the Ser368 hydroxyl oxygen on carbonyl carbon of P1. The MD simulations showed that even for the flexible short peptide studied here, only the MHV-1 and MHV-3 strains produced complexes that satisfy these criteria, in agreement with the cleavage assays where only these two strains were cleaved among the five tested. Interestingly, we have found that substitution of the canonical Arg in the P1 substrate position by Ser or Thr (in the MHV-2 and MHV-S strains, respectively) disrupts salt-bridges that form in the active site, which leads to distorted active site conformations in which the furin catalytic triad residues that are far from their positions needed for enzymatic activity. This result aligns with both the sequence-based ProP prediction and the findings from the cleavage assays for the MHV-2 and MHV-S strains. For the MHV-A59 strain in which Arg in the P2 substrate position was replaced by His, the ProP algorithm suggested a high possibility for furin cleavage– a prediction that was not supported by the results of the cleavage assays. In the MD simulations, we found that the P2 histidine in MHV-A59 does create nonbonded interactions with the residues that form the canonical P2 Arg binding site, but these interactions are not sufficient to sustain function because this P2 residue does not properly orient the His194 of the furin catalytic triad to act as the required proton switch. Consequently, the active site conformation of the furin/MHV-A59 complex does not comply with the structural requirements for the cleavage initiation despite the assumed similarity of a protonated His with the positively charged Arg site chain. This result underscores the need to consider the structural aspects of both the active site and the putative substrate for furin-mediated cleavage. Notably, this implies that expanding these mechanistic findings to predicting the cleavability of the S1/S2 cleavage sites in spike glycoproteins would require additional assessment of the conformational dynamics of the full-length loop segments carrying the cleavage site in all-atom models of the spike protein.

### Limitations of this study

As outlined in Jaimes et al. (22, 23) the short peptides used in this study do not reflect the tertiary structure of the spike glycoprotein in vivo and hence the conformations they may present to the furin active site, along with the relative orientation of the peptide in the furin binding pocket.

## ACKNOWLEDGEMENTS

We thank Dr. Avery August and Dr. Melanie Ragin for their support and mentorship of D.S. within the Howard Hughes Medical Institute Cornell University Research Transfer Program. A.C. is supported by grant T32EB023860 from the National Institute of Biomedical and Bioengineering. Work on coronavirus entry in the Whittaker lab is funded in part by the National Institute of Health research grant R01AI35270. Access to computational resources allocated by the COVID-19 High Performance Computing Consortium at the Oak Ridge Leadership Computing Facility, which is a DOE Office of Science User Facility supported under Contract DE-AC05-00OR22725, and the AiMOS supercomputer at the Center for Computational Innovations (CCI) of Rensselaer Polytechnic Institute (RPI), is gratefully acknowledged together with the in-house computational resources of the David A. Cofrin Center for Biomedical Information in the Institute for Computational Biomedicine at Weill Cornell Medical College.

## MATERIALS AND METHODS

### Prediction of furin cleavage via ProP Software

First, to predict whether furin cleavage occurs at the S1/S2 domain of the S protein, we use ProP computer software as discussed in Duckert et al. (2004). According to Duckert et al. (2004) a score greater than 0.5 suggests a sufficient prediction of a furin cleavage site. Within the ProP software, we recorded the series of amino acid residues of the S protein for coronaviruses under investigation, and then received a score.

### Accession numbers

Accession numbers of spike protein sequences used in this study are as follows: SARS-CoV-1 (YP_009825051.1), SARS-CoV-2 (YP_009724390.1), MERS-CoV (NC_01984.3), HCoV-OC43 (NC_006213.1), HCoV-HKU1 (NC_006577), FCoV-UU4 (ACT10887.1), MHV-S (AFD97607), MHV-1 (ACN89742.1), MHV-2 (AF201929_4), MHV-3 (QSJ02954.1), MHV-JHM (YP_209233), MHV-A59 (YP_009824982)

### Fluorogenic Peptide Cleavage Assay

The fluorogenic peptide cleavage assay, along with furin and trypsin buffer dilutions, were conducted essentially as in Jaimes et al. (2017). First, we designed and ordered linear peptides that mimic the S1/S2 domain of the S glycoprotein for selected coronaviruses (Biomatik, Kitchener, Ontario). Information about the series of amino acid residues within the S1/S2 domain were obtained from NCBI, by searching for the accession numbers/complete genomes of MHV-1, MHV-2, MHV3/JHM, MHV-S, and MHVA59, followed by a multiple sequence alignment. Each of the peptide mimics used contained a fluorescence resonance energy transfer (FRET) pair, 7-methoxycoumarin-4-yl-acetly (MCA) and 2,4-dinitrophenol (DNP), on the N-terminus and C-terminus ends of the peptide. MCA is a light-sensitive molecule where fluoresence is quenched in the presence of DNP. However, if a protease recognizes a site on the peptide, it is able to cleave it, and upon cleaving DNP no longer quenches MCA’s light and that signal is detected by a fluorescent plate reader. From this signal, we acquire information about the processivity of the protease, and can do a comparison between peptides within the same biological replicate. Three biological replicates were completed for each of the furin buffer pH values (four total) under investigation.

In a 96-well plate, each well contained one of our selected S1/S2 peptides mixed in a solution of diluted protease with its buffer. The proteases used in this experiment include the prototypical proprotein convertase furin (New England BioLabs) and the prototype serine endopeptidase trypsin (New England BioLabs), placed within a fluorescence plate reader (Spectra max Gemini XPS, Molecular Devices) to measure cleavage, and then monitored by an increase in fluorescence based on the separation of FRET pair for one hour. Similar to Jaimes et al. (22), data from the fluorescent plate reader was saved as a.txt file and analyzed within Excel (Microsoft). For each biological replicate, we averaged three technical replicates, to obtain a graph where relative fluorescent units were plotted on the y-axis and time on the x-axis. From the slope of the graph, we obtained the Vmax of protease, and compared the processivity of furin cleavage for each peptide mimic. The three biological replicates were averaged to obtain the final Vmax.

### Preparation of the molecular systems

The initial models for the MD simulations of the protein-peptide complexes of furin with each of the five peptide strands are based on the X-Ray structure of furin in complex with substrate-like inhibitor 4-aminomethyl-phenylacetyl-canavanine-Tle-Arg-Amba (PDB ID 6YD2) (10). In building the complexes, the peptides were positioned with the putative P1 to P4 residues aligned to those of the inhibitor molecule observed in the X-Ray structure. Each furin-peptide complex was solvated in an all-atom environment of a TIP3 water solution of 0.15 M sodium chloride, contained in rectangular boxes with the minimum and maximum box padding set to 15 Å.

### Equilibration of MD systems

Each system was equilibrated in a three-step equilibration procedure in which the system was simulated for 30ns per each step, for a 90ns total equilibration time per complex. In step 1, harmonic constraints were imposed on the backbone of furin and the substrates, with harmonic constant 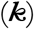 values of 5 and 25 kcal/(mol*Å^2^), respectively. These constraints were gradually released in the following steps. The constrains on the furin backbone were released in step 2, while constraints were loosened on the backbone atoms of residues in positions P5 to P1’ of the peptide, with a 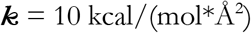. All the constrains in the systems were released in step 3. For the production stage, 5 replicas were constructed for each of the five furin-peptide complexes and MD simulations of each replica were run for 150 ns for a total of 750 ns-long sampling per complex.

### Parameters of MD simulations

All the simulations were done with NAMD2.13 software (24) and the CHARMM36 force field (25). The temperature (310 K) and pressure (1 bar) control was implemented with Langevin Thermostat (*damping coefficient* = 5/ps) and Langevin Piston (*period* = 100 fs, *damping time constant* = 50 fs). Long-range electrostatic interactions were treated with Particle Mesh Ewald (PME) approximation. Integration timestep was set to 2 fs, cutoff and switching distances for non-bonded interactions were set to 10 Å and 13.5 Å respectively. All bonds involving hydrogen were fixed (*rigidBond* = all).

### Analysis of the MD trajectories

Analyses of the MD trajectory data for each complex was carried out to reveal the resulting structural details of the catalytic sites and the behavior of the water surrounding these areas. To quantify water accessibility of catalytic triad region of the peptide binding site, we evaluated the number of MD frames in which at least one water molecule was located within 2.8 Å of both hydroxyl oxygen of Ser368 and carbonyl carbon of P1 residue with *compute_neighbors* tool of MDtraj python package (26).

---

**Figure S1.**
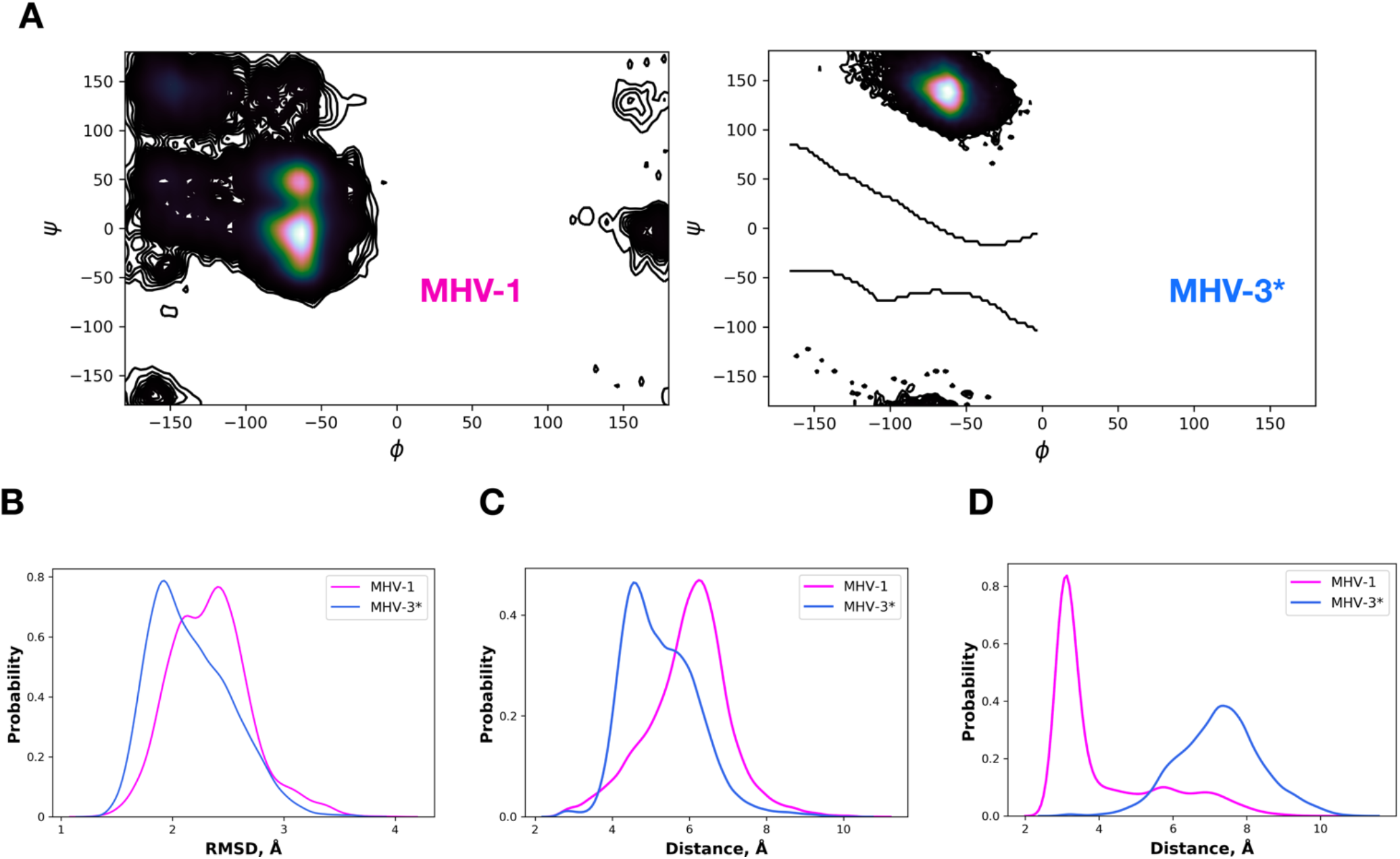
The effect of His and Arg residues in the P5 position of the substrate on the geometry and dynamics of the substrate binding site recorded in the MD simulations of furin complexes with MHV-1 and MHV-3 strains peptides. A) Distribution of the backbone dihedrals between P5 and P6 residues of MHV-1 and MHV-3 in their complexes with furin. B) Profile of the Root Means Square Deviation (RMSD) values for the backbone atoms of MHV-1 and MHV-3 peptides in MD simulations of their complexes with furin. C-D) Profile of the Minimum distances measured between heavy atoms of the substrate P5 residues in MHV-1 and MHV-3 and the side chains of either Glu230 or Glu257 (C) and Val231 (D) from MD simulations of furin in complex with MHV-1 or MHV-3 strains

## Notes

### Competing Interest Statement

The authors have declared no competing interest.

